# Genome-wide DNA methylation profiling of the zebrafish forebrain

**DOI:** 10.64898/2026.03.20.713102

**Authors:** Pol Sorigue, Maeva Pinget, João Costa, Magda C. Teles, Rui F. Oliveira

**Affiliations:** GIMM - Gulbenkian Institute for Molecular Medicine, Rua Quinta Grande 6, 2780-156 Oeiras, Portugal; ISPA - University Institute for Psychological, Social and Life Sciences, Rua do Jardim do Tabaco 34, 1149-041 Lisbon, Portugal

## Abstract

The zebrafish forebrain is a central hub for cognitive and social behaviors, yet a comprehensive, region-specific DNA methylation reference for this tissue is not available. To address this gap, we generated a genome-wide methylation map of the adult zebrafish forebrain using Oxford Nanopore Technologies (ONT) long-read sequencing. Forebrains from six individuals were profiled, enabling direct detection of multiple DNA base modifications at single-base resolution. The resulting dataset includes CpG-associated 5-methylcytosine (5mC) and 5-hydroxymethylcytosine (5hmC), as well as non-CpG 5mC and N6-methyladenine (6mA), with 96.8% genome coverage. CpG methylation was widespread, with 64.2% of CpG sites classified as highly 5mC-modified. In contrast, 5hmC and non-CpG methylation were markedly less abundant, and 6mA was detected at very low levels. CpG islands showed a bimodal distribution of methylation, promoter regions displayed elevated variability compared to gene bodies, and CC-context non-CpG methylation exhibited significant strand asymmetry. This resource provides a high-resolution epigenomic baseline for the adult zebrafish forebrain.

## Background & Summary

The forebrain is a principal center for higher-order neural processing, integrating sensory information with internal physiological states to generate context-appropriate behavioral responses. It plays a central role in cognition, learning and memory, decision-making, and the regulation of complex social behaviors ^1^. In teleosts, forebrain regions homologous to the mammalian telencephalon and diencephalon are critically involved in social behavior, emotional processing, and responses to environmental stimuli ^2,3^. These regions form interconnected circuits that evaluate social cues, assess environmental challenges, and coordinate adaptive behavioral outputs ^4,5^.

Importantly, the functional plasticity of the forebrain is not determined solely by neural circuitry but is dynamically shaped by molecular mechanisms that regulate gene expression ^6^. Among these, epigenetic processes provide a key interface between environmental experience and long-term changes in brain function ^7^. DNA methylation, in particular, has been strongly associated with neural plasticity, learning, and the emergence of distinct behavioral phenotypes across vertebrates ^8^. In mammals, early-life experiences can induce persistent alterations in DNA methylation patterns within forebrain structures, thereby influencing stress reactivity, cognition, and social behavior later in life ^9,10^. Moreover, locus-specific methylation differences have been linked to individual variation in behavioral traits ^11–13^. Similarly, in teleosts, socially relevant environmental changes, such as shifts in social status ^14,15^ or territorial intrusion ^16^ are associated with genome-wide epigenetic reprogramming in the brain. Environmentally induced epigenetic modifications have also been shown to shape social phenotypes and behavioral strategies ^17^.

Among all epigenetic modifications, DNA methylation is one of the most extensively studied. It involves the addition of methyl groups to cytosine or adenine residues and can modulate gene expression without altering the underlying DNA sequence. In vertebrates, DNA methylation predominantly occurs as 5-methylcytosine (5mC) at CpG dinucleotides. CpG-associated 5mC is linked to transcriptional repression, genomic imprinting, X-chromosome inactivation, and transposable element silencing ^18,19^. Promoter and CpG island methylation is generally associated with stable gene silencing, whereas gene body methylation is often found in actively transcribed genes and may enhance transcriptional fidelity by limiting spurious initiation ^20,21^. An oxidized derivative of 5mC, 5-hydroxymethylcytosine (5hmC), is particularly abundant in the central nervous system, where it accumulates in post-mitotic neurons and is enriched within gene bodies of actively expressed neuronal genes ^22,23^. Beyond CpG methylation, 5mC also occurs at non-CpG sites (CpH, where H = A, C, or T), which are relatively rare in most somatic tissues but enriched in the nervous system ^24^. In addition, N6-methyladenine (6mA) has been reported in vertebrate genomes, although its abundance and functional relevance remain under investigation ^25,26^.

Advances in epigenomic profiling technologies have enabled genome-wide detection of DNA base modifications at single-base resolution. The first base-resolution zebrafish brain methylome was obtained using Reduced Representation Bisulfite Sequencing ^27^. Although bisulfite-based sequencing approaches remain widely used for mapping 5mC modifications, single-molecule sequencing platforms, including those developed by Oxford Nanopore Technologies (ONT), permit the direct detection of multiple DNA modifications without chemical conversion or amplification ^28,29^. A recent study of the zebrafish kidney methylome reported biases associated with whole-genome bisulfite sequencing that were not observed when methylation was profiled using nanopore-based approaches ^30^. Furthermore, nanopore sequencing enables simultaneous profiling of CpG and non-CpG 5mC, 5hmC, as well as 6mA. Nevertheless, detection of low-abundance modifications remains technically challenging, and signal discrimination for marks such as 6mA and non-CpG methylation in vertebrate genomes requires careful validation ^31^.

Despite increasing recognition of DNA methylation as a key regulator of brain function, high-resolution epigenomic maps of the zebrafish brain remain limited. The zebrafish (*Danio rerio*) offers a powerful vertebrate model for investigating the molecular basis of neural and behavioral plasticity. As a teleost with conserved forebrain organization and substantial genetic homology to mammals, it combines experimental accessibility, well-characterized social behaviors, and a high-quality, extensively annotated genome ^32^. Here, we present a comprehensive long-read methylome dataset of the adult zebrafish forebrain generated using Oxford Nanopore sequencing, comprising genome-wide maps of CpG-associated 5mC and 5hmC, as well as non-CpG 5mC and 6mA. Given that vertebrate neural tissues exhibit distinctive and highly specialized DNA methylation landscapes, this dataset provides a region-specific epigenomic reference that will facilitate comparative, developmental, and functional analyses of vertebrate brain methylation dynamics.

## Methods

### Animals and housing conditions

All animals used in this experiment were wild-type zebrafish (TL background) bred and maintained at the Gulbenkian Institute for Molecular Medicine (GIMM, Oeiras, Portugal). Fish were placed in a Tecniplast recirculating life support system at 5 days post-fertilization (dpf) and maintained in a polyculture of rotifers and algae until 8 dpf, after which the feeding regime was initiated. They were fed twice daily: rotifers in the morning and commercial food flakes (Gemma) in the afternoon. All animals were kept at 28 °C under a 14:10 h light-dark photoperiod, with a pH of 7.0 and conductivity of 1200 μS/cm. All experiments were conducted in accordance with the standard operating procedures of the Gulbenkian Institute for Molecular Medicine Ethics Committee. No permit number was required for this experiment, as the animals were maintained under standard housing conditions and no experimental procedures were performed. All procedures were carried out in accordance with relevant guidelines and regulations, namely in compliance with the Portuguese law, DL 113/2013, which transposes the Directive 2010/63/UE of the European Parliament and Council, for the protection of animals used for scientific experimental purposes. All methods are reported in accordance with ARRIVE guidelines ^33^.

### Sample collection

A total of six fish (2 females and 4 males), aged 4 months ± 1 week, were randomly selected from three different wild-type tanks. Fish were euthanized with an overdose of tricaine solution (MS-222, Pharmaq; 1200 mg/L), and the forebrain (telencephalon and diencephalon) was dissected under a stereomicroscope (Nikon, SMZ 745). After dissection, the forebrain tissue was collected and placed in 1.5 mL tubes containing 100 µL of HMW gDNA Tissue Lysis Buffer from the Monarch High Molecular Weight (HMW) DNA Extraction Kit for Tissue (New England Biolabs; NEB #T3060). Samples were immediately placed on dry ice and stored at −80 °C until further use. Biometric data were also collected, including standard length (measured from the tip of the snout to the end of the caudal peduncle) 2.53 ±0.11, and body weight (g) 0.30 ± 0.02.

### DNA extraction

Genomic DNA was extracted using the Monarch® High Molecular Weight (HMW) DNA Extraction Kit from Tissue (New England Biolabs, NEB #T3060), following the adapted manufacturer’s instructions to accommodate very low tissue inputs (< 2–5 ng of starting material). Briefly, brain tissue was mechanically dissociated using a sterile pestle. The kit uses a glass bead–based purification method optimized for the isolation of extremely high–molecular–weight DNA, combining gentle cell lysis with controlled DNA fragmentation and subsequent precipitation of DNA onto the surface of the large glass beads. Protocol modifications were implemented to minimize mechanical agitation and preserve DNA integrity. Under these conditions, extracted DNA fragment sizes typically ranged from 50–250 kb using the standard protocol, with the potential to reach megabase-scale fragments when the lowest agitation speeds were employed. Purified DNA was eluted into low-binding tubes (Eppendorf) and stored at 4 °C until library preparation.

### Library preparation and Nanopore sequencing

Sequencing libraries were prepared using the Rapid Barcoding Kit 24 v14 (SQK-RBK114-24; Oxford Nanopore Technologies) according to the manufacturer’s instructions. Each sequencing run included six individuals, resulting in 4 independent library preparation and sequencing runs. Sequencing was performed on a PromethION platform using FLO-PRO114M flow cells and MinKNOW software (version 25.05.14).

### Basecalling, alignment, and quality control

Raw Nanopore sequencing data (FAST5 format) were basecalled using Dorado (version 1.3.1; https://github.com/nanoporetech/dorado) with the dna_r10.4.1_e8.2_400bps_sup@v5.0.0 model. Basecalling included detection of DNA base modifications, specifically 5-methylcytosine (5mC), 5-hydroxymethylcytosine (5hmC), and N6-methyladenine (6mA). Reads with a quality score below 10 were automatically filtered out by MinKNOW software. Adapter sequences were trimmed during basecalling. Output files were generated in modBAM format, which extends the BAM format by including per-base modification probability annotations.

Basecalled reads were aligned to the zebrafish reference genome assembly GRCz12tu using Dorado with default alignment parameters. Only high-confidence primary alignments were retained by removing reads with mapping quality (MAPQ) < 10, secondary and supplementary alignments, unmapped reads, and reads shorter than 200 bp. Filtered BAM files were subsequently sorted and indexed using samtools (version 1.21; https://github.com/samtools/samtools).

Genome recovery rate was calculated for each replicate and across replicates using the samtools *coverage* command to determine the proportion of the genome covered by at least one read.

### Modification quantification

Read-level modification statistics were obtained using modkit (version 0.5.0; https://nanoporetech.github.io/modkit) *summary* command. We used a confidence threshold of 0.8 to call the presence of a modification or a canonical base-pair. To obtain read-level statistics, a random subset of 100,000 reads per replicate was sampled.

### Methylation profiling and dataset generation

The modBAM files were converted to bedMethyl format using modkit with the *pileup* command. The bedMethyl format enables representation of per-base methylation levels by reporting genomic coordinates, strand information, read coverage, and modification frequencies. A confidence threshold of 0.8 was applied uniformly to call the presence or absence of a modification at each base.

Three bedMethyl datasets were generated:

1. A comprehensive dataset containing all detected base modifications with strand-specific information.
2. A CpG-restricted dataset in which calls were limited to CpG dinucleotides and aggregated across strands.
3. A CH-restricted dataset (CA, CC, CT motifs) retaining strand-specific information.

Strand aggregation for CpG sites was performed because CpG methylation is typically symmetric across complementary strands, allowing reads from both strands to contribute to a single genomic coordinate. The mitochondrial chromosome was excluded from all downstream analyses. For downstream processing, bedMethyl tables were reduced to include genomic position (0-based coordinate of the modified base), read coverage, methylation fraction (number of modification calls divided by the total coverage at that position), and strand.

### Motif coverage and modification assessment

For all sequence motifs analyzed (CG, CC, CT, CA, and A), motif occurrence was evaluated on a per-sample basis using the modkit *motif evaluate* command. This command computes the number of motif occurrences associated with each modification type and categorizes them as low, mid, or high modified based on the fraction of reads supporting a modification. Default modkit thresholds were used to define modification categories: motif instances with a modified-read fraction < 0.2 were classified as low modified, whereas those with a modified-read fraction >0.6 were classified as high modified; remaining sites were classified as mid modified. For each motif–modification combination, we calculated the proportion of highly modified sites as the number of sites with a modified-read fraction > 0.6 divided by the total number of covered motif occurrences. Only positions supported by ≥5x read depth were considered in the analysis.

### Genomic distribution of DNA modifications

Genomic distribution analyses were performed using the following region definitions:

1. **Genomic bins**: The zebrafish genome was partitioned into unsupervised non-overlapping 50 kb bins using bedtools (version 2.31.1; https://github.com/arq5x/bedtools2) *makewindows* command, excluding the mitochondrial chromosome.
2. **CpG islands**: Annotated CpG islands for the zebrafish GRCz12tu genome assembly were downloaded from the UCSC Genome Browser (hgTables) on 22 October 2025 (https://genome.ucsc.edu/).
3. **Promoters:** Promoter regions were defined as the genomic interval spanning 2,000 bp upstream of the transcription start site (TSS) to the TSS for all annotated transcripts, based on the NCBI GTF annotation for the zebrafish GRCz12tu genome assembly.
4. **Gene bodies**: Gene bodies were defined as the region from the TSS to the transcription end site (TES) for all annotated genes in the NCBI GTF file.

Only positions with ≥5x read coverage were included. Mean methylation levels per region were calculated using bedtools *map* command as the average of per-site methylation fractions across all covered positions within the region. To ensure robust estimates, regions containing fewer than a minimum number of motif occurrences were excluded from the analysis (100 for genomic bins, 5 for CG islands, 10 for promoters, 20 for gene bodies). Regions were analyzed independently, and overlapping regions were not mutually exclusive.

For CpG motifs in all region types, and genomic bins, strand information was ignored. In contrast, for promoter and gene body regions, overlaps were restricted to cases where the modification position and the annotated region were on the same strand. Strand symmetry was further assessed for non-palindromic motifs by analyzing positive and negative strands separately using strand-specific datasets. Due to the extremely low abundance of 6mA modifications, this modification type was excluded from downstream regional analyses.

Replicate-averaged methylation values were computed by first averaging methylation per region within each replicate, followed by aggregation across replicates to generate a single value per region. This approach reduces technical variability while ensuring equal regional contribution.

Strand-specific differences were assessed using Welch’s two-sample t-tests based on replicate-averaged values. P-values were adjusted for multiple testing using the Benjamini–Hochberg false discovery rate (FDR) procedure, applied separately for each motif.

### Forebrain-specific methylation analysis

To contextualize CpG methylation levels, we compared our dataset with a previously published zebrafish whole-brain methylome ^27^ generated using reduced representation bisulfite sequencing (RRBS). Raw FASTQ files were downloaded and aligned to the zebrafish GRCz12tu genome assembly using Bismark (version 0.25.1) with the *bismark* command. Methylation calls were generated using the *bismark_methylation_extractor* command, and strand-combined methylation calls were obtained using the *--comprehensive* flag.

Because RRBS-derived methylation signals reflect both 5-methylcytosine (5mC) and 5-hydroxymethylcytosine (5hmC), counts for both modification types were summed at each CpG position in our Nanopore-based forebrain dataset. Mean methylation levels were then computed using the bedtools *map* command, following the same procedure described above, for both whole-brain and forebrain datasets. This yielded a dataset of CpG island locations with corresponding methylation levels for each condition. To ensure robust estimates, CpG islands with a mean read depth below 10 or fewer than 20 covered CpG sites were excluded.

Differences between forebrain and whole-brain datasets were calculated as delta methylation (forebrain − whole brain). CpG islands were classified as hyper- or hypomethylated using an absolute delta threshold of 0.15. Gene bodies and promoters overlapping CpG islands exceeding this threshold were identified for annotation purposes.

## Data records

Raw sequencing reads and methylation BAM files for all samples have been deposited in the European Nucleotide Archive under study accession PRJEB108899. Processed methylation matrices separated by motif and modification type, together with region-level methylation files, are available via ArrayExpress under accession E-MTAB-16780.

Methylation matrices use 0-based genomic coordinates and include genomic position, coverage, and the number of methylation calls. For non-CpG motifs, strand information is also provided. Region-level methylation files are likewise reported using 0-based coordinates and aggregate all region types, motifs, and replicates into a single file through clearly labelled metadata columns.

Additional intermediate analysis files are available from the corresponding author upon reasonable request.

## Technical validation

### Quality Control

BAM files from each replicate were filtered to retain high-confidence alignments by applying the following criteria: mapping quality (MAPQ) > 10, read length > 200 bp, and exclusion of supplementary, secondary, and unmapped reads (Figure 1). After filtering, the dataset comprised 25,169,629 aligned reads spanning 1,402,091,628 bases of the reference genome. This provided 96.78% coverage of the reference genome at a minimum depth of 1×. The mean base quality score across retained reads was 41.3, and the mean mapping quality was 57.95. Detailed sequencing and alignment statistics for each replicate, including read counts before and after filtering, coverage metrics, and quality score distributions, are reported in Supplementary Table 1.

**Figure 1.**
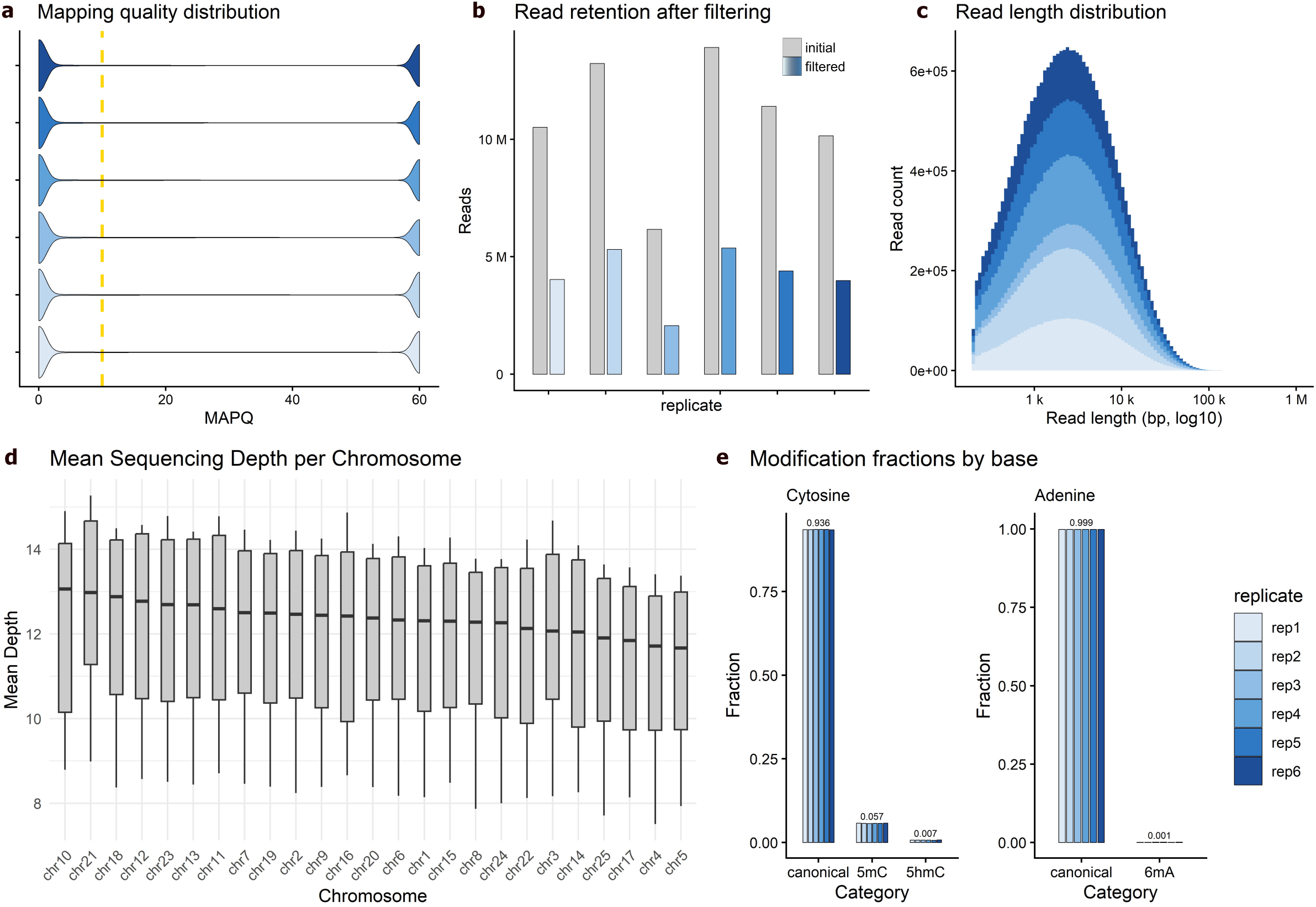
Sequencing quality metrics and global modification statistics for Nanopore forebrain methylomes: **(a)** Distribution of mapping quality (MAPQ) scores for aligned reads across all replicates. The dashed yellow line indicates the MAPQ threshold (MAPQ ≥ 10) used for read filtering. **(b)** Number of reads retained before (grey) and after (blue) filtering for each replicate, following removal of low-quality, secondary, supplementary, unmapped, and short (<200 bp) reads. **(c)** Read length distributions for filtered reads across replicates, shown on a logarithmic x-axis. **(d)** Distribution of mean sequencing depth per chromosome across replicates, illustrating uniform genome-wide coverage after filtering. Boxplots show median, interquartile range, and range of mean depth values per chromosome. **(e)** Fraction of canonical and modified base calls for cytosine (left) and adenine (right). Cytosines were classified as canonical, 5-methylcytosine (5mC), or 5-hydroxymethylcytosine (5hmC), while adenines were classified as canonical or N6-methyladenine (6mA). Fractions represent read-level modification calls pooled per replicate. Numeric labels indicate median value of all replicates.

### Modification calling summary

We quantified the number of base calls classified as canonical or modified across all replicates. Cytosines were categorized as canonical, 5-methylcytosine (5mC), or 5-hydroxymethylcytosine (5hmC), while adenines were classified as canonical or N6-methyladenine (6mA). Across replicates, cytosine calls were predominantly canonical (93.6%), with 5.7% classified as 5mC-modified and 0.7% as 5hmC-modified. Adenine modification was rare, occurring at a frequency of approximately one modified adenine per 1,000 adenines (0.1%; Figure 1). Importantly, these values represent individual modification calls on independent sequencing reads; integration across genomic coordinates is addressed in subsequent analyses. Detailed values and standard errors can be consulted in Supplementary Table 1.

### Motif-level distribution and modification frequencies

We quantified cytosine modifications occurring at CpG and non-CpG genomic sites, as well as adenine modifications. A genomic position was classified as modified if the fraction of modified reads at that position was ≥0.6, as computed using the modkit *motif evaluate* default filtering threshold. Across replicates, 64.2 ± 0.45% (mean ± s.d.) of genomic CpG motifs were classified as 5mC-modified. In contrast, 0.41 ± 0.01% of CpG motifs were classified as 5hmC-modified. Modification frequencies at non-CpG cytosine motifs (CA, CC, CT) were substantially lower (Figure 2). Adenine methylation (6mA) was detected at extremely low levels, corresponding to approximately five modified sites per 10⁸ genomic adenines (Figure 2; Supplementary Table 2). Detailed values and standard errors can be consulted in Supplementary Table 2.

**Figure 2.**
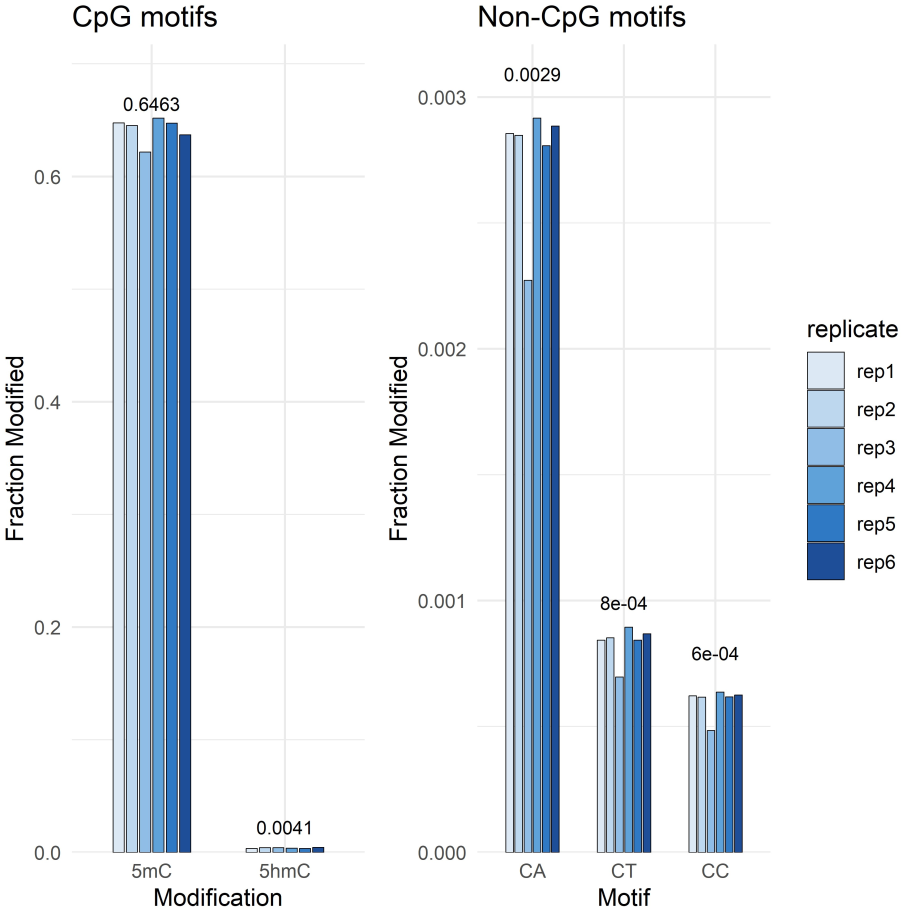
Genome-wide distribution of cytosine methylation across CpG and non-CpG motifs: Fraction of modified cytosines across replicates for CpG (left) and non-CpG (right) motifs. For CpG sites, cytosines were classified as 5-methylcytosine (5mC) or 5-hydroxymethylcytosine (5hmC). For non-CpG motifs, 5mC levels are shown separately for CA, CT, and CC contexts. Bars represent per-replicate fractions of modified calls relative to the total number of motif occurrences. Numeric labels indicate the median fraction across replicates. 5hmC modifications at non-CpG cytosine motifs are not reported because their frequencies are extremely low and considered negligible. However, these values can be consulted in Supplementary Table 2. CpG sites show high levels of 5mC and low 5hmC, whereas non-CpG methylation is present at substantially lower frequencies across all contexts.

### Genomic distribution of CpG motif methylation

CpG-associated 5mC methylation levels in gene bodies were comparable to those observed in genomic bins (gene bodies: median 0.65, IQR 0.60–0.71; genomic bins: median 0.66, IQR 0.56–0.72). In contrast, promoter regions exhibited greater heterogeneity and relatively lower median methylation levels (median 0.61, IQR 0.36–0.79). CpG islands displayed a bimodal distribution of 5mC methylation values, with a major proportion of islands showing either low or high methylation levels (Figure 3). CpG-associated 5hmC methylation levels were consistently lower than 5mC across all genomic features analyzed. Median 5hmC values were below 0.1 in all region types analyzed, with narrower interquartile ranges. Detailed values can be consulted in Supplementary Table 3.

**Figure 3.**
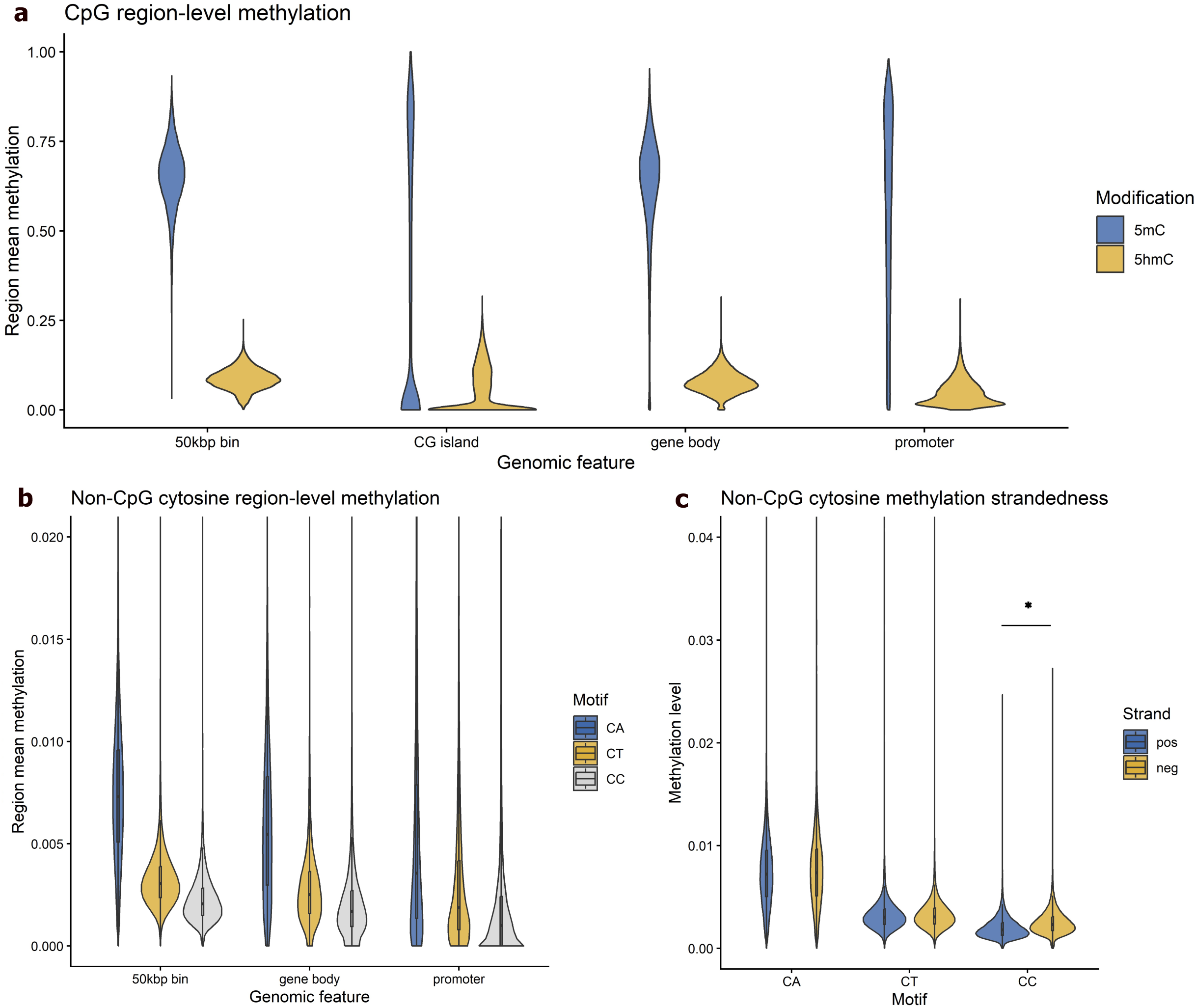
Regional distribution and strand specificity of cytosine methylation: **(a)** Distribution of region-level mean methylation values for CpG-associated cytosine modifications across unsupervised 50 kbp genomic bins, CpG islands, gene bodies, and promoter regions. Violin plots show the distribution of methylation values per-region for 5-methylcytosine (5mC) and 5-hydroxymethylcytosine (5hmC). **(b)** Region-level mean methylation of non-CpG cytosines stratified by motif (CA, CT, and CC) across 50 kbp bins, gene bodies, and promoters. Non-CpG methylation levels are substantially lower than CpG methylation and vary by motif and genomic context. **(c)** Strand-specific methylation levels for non-CpG motifs across unsupervised 50 kbp bins. Positive and negative strands were analyzed independently to assess strand asymmetry. An asterisk indicates motifs showing a significant difference between strands after multiple testing correction (Welch’s t-test, Benjamini–Hochberg FDR). * adjusted p-value = 4.2x10^-5^

### Genomic distribution of non-CpG motif methylation

Region-level 5mC methylation at non-CpG cytosines (CA, CC, and CT motifs) was low across all genomic features examined (Figure 3). Among the three motifs, CA sites exhibited the highest methylation levels, followed by CT and CC. This ranking was consistent across 50 kbp genomic bins, gene bodies, and promoter regions. In 50 kbp bins, median methylation levels were 0.0073 (IQR 0.0051–0.0096) for CA motifs, 0.0031 (IQR 0.0024–0.0039) for CT motifs, and 0.0021 (IQR 0.0015–0.0028) for CC motifs. Gene bodies showed slightly lower methylation levels overall, while promoters displayed the lowest median values across all motifs, accompanied by broader interquartile ranges, particularly for CA and CT motifs (Figure 3). Detailed values can be consulted in Supplementary Table 3.

Strand symmetry of non-CpG 5mC methylation was evaluated at the regional level. CC motifs showed significant strand asymmetry across all region types (genomic bins, promoters, and gene bodies), and across all chromosomes (p < 0.005). In contrast, CA and CT motifs did not exhibit significant strand asymmetry in any region type or chromosome (Figure 3), with the exception of CT motifs on chromosome 8 (p = 0.004). Detailed statistical results are provided in Supplementary Table 4.

### Forebrain-specific methylation analysis

To assess the technical reliability of the Nanopore-based forebrain methylome, CpG island methylation levels were compared with a previously published zebrafish whole-brain dataset generated using reduced representation bisulfite sequencing ^27^. CpG islands were retained if they had ≥20 covered CpG sites and a mean read coverage >10, resulting in 2,778 islands for analysis.

Methylation levels between the two datasets were highly concordant, with Pearson and Spearman correlations of r = 0.98 and ρ = 0.91, respectively (Figure 4). These strong correlations indicate that the Nanopore-based methylation estimates accurately capture CpG island methylation patterns, despite differences in tissue specificity and sequencing technology.

**Figure 4.**
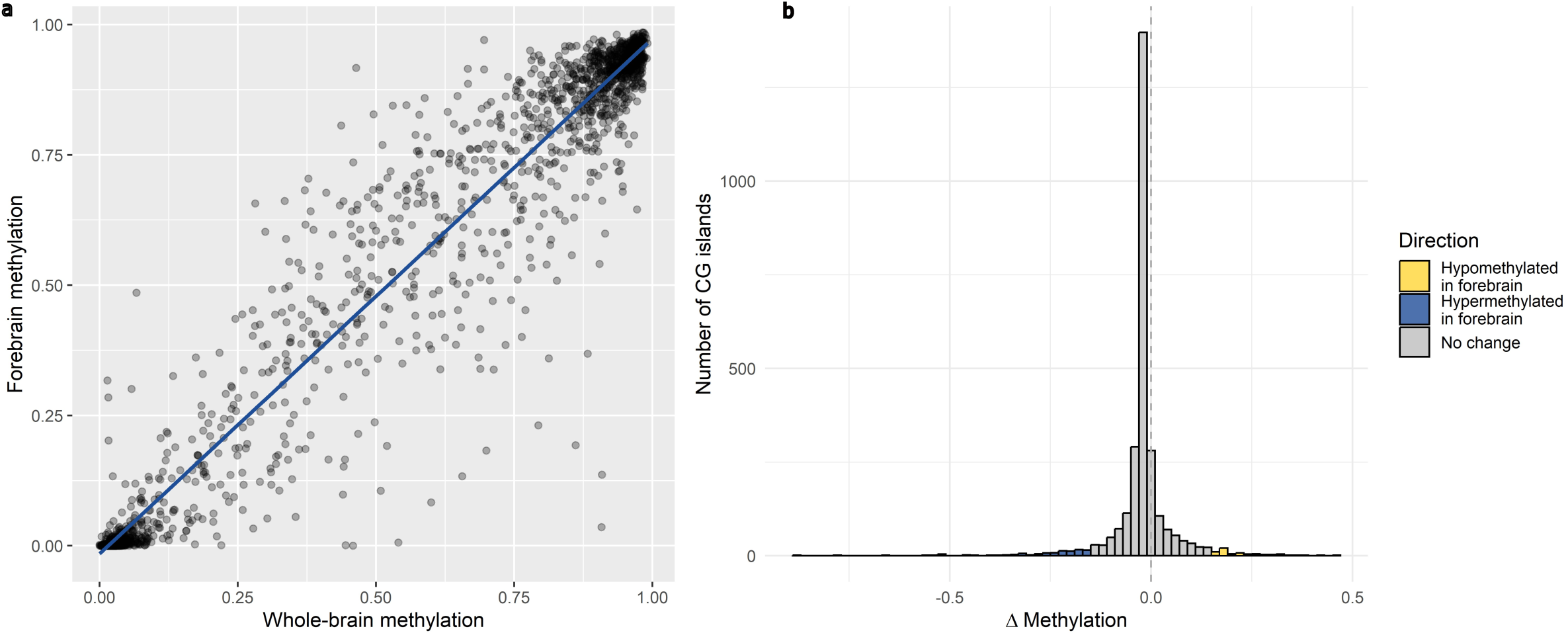
Comparison of CpG island methylation between whole brain and forebrain: **(a)** Scatter plot of mean CpG island methylation levels in whole-brain versus forebrain datasets (n = 2,778 CpG islands). Each point represents one CpG island passing coverage and CpG count filters. The solid line shows the linear regression fit. Methylation levels were highly concordant between datasets (Pearson r = 0.98; Spearman ρ = 0.91). **(b)** Distribution of differential CpG island methylation (Δ methylation = forebrain − whole brain). CpG islands were classified as hypomethylated (yellow) or hypermethylated (blue) in the forebrain using an absolute Δ methylation threshold of 0.15; CpG islands not exceeding this threshold are shown in gray. The dashed vertical line indicates no methylation difference.

A small subset of CpG islands showed substantial differences between forebrain and whole-brain methylation, defined as an absolute delta methylation (forebrain − whole brain) > 0.15. This threshold identified 103 hypomethylated and 66 hypermethylated CpG islands in the forebrain, while 2,609 CpG islands remained largely unchanged (Figure 4; Supplementary Table 5).

Differentially methylated CpG islands between forebrain and whole-brain datasets overlapped a broad set of genes, including those associated with neuronal development, transcriptional regulation, signal transduction, and synaptic function. Forebrain-hypomethylated islands included regions overlapping neurodevelopmental regulators and transcription factors such as *pax3a*, *neurod6a*, *npas3*, *foxp4*, *irx2a*, *lhx3*, *barhl2*, and *prox2*, as well as genes involved in synaptic signaling and neuronal connectivity, including *nlgn2b*, *nlgn4xb*, *reln*, multiple *protocadherin* genes, and *kirrel1a*/*kirrel3a*. Several ion channels and signaling molecules, including *kcnj6*, *kcnn1a*, *kcnk15*, *pde3a*, and *rgs22*, were also associated with forebrain-specific hypomethylation. Conversely, hypermethylated CpG islands in the forebrain overlapped genes implicated in cell signaling, cytoskeletal organization, transcriptional control, and synaptic structure, including *znf462*, *zbtb49*, *cxxc5a*, *med1*, *pbx1b*, *nfixa*, *zeb2b*, *celsr2*, *plxnb2b*, *talin2b*, and *nbeal1*.

Overall, the high concordance with an independent RRBS dataset provides strong technical validation for the Nanopore-based forebrain methylome and supports the reliability of base modification calls across CpG islands.

## Data availability

All raw sequencing data have been deposited in the European Nucleotide Archive (ENA) under the study accession number PRJEB108899. Methylation matrices and region-level methylation files have been deposited in ArrayExpress under accession E-MTAB-16780.

## Code availability

All software tools used in this study are publicly available, and the corresponding parameter settings are described in the Methods section. Where parameters are not explicitly stated, default settings recommended by the respective software developers were applied. Custom scripts developed for data processing and analysis are available at https://github.com/psorigue/Zebrafish-Forebrain-Methylome, with a permanently archived version accessible via Zenodo (DOI:10.5281/zenodo.18985815).

## Acknowledgements

We would like to thank the Genomics Facility of the Gulbenkian Institute for Molecular Medicine (GIMM) for their sequencing support and technical assistance. We also thank the Aquatic Facility of the Gulbenkian Institute for Molecular Medicine for providing zebrafish infrastructure and animal care.

## Author Contributions

Pol Sorigue: Methodology, Software, Validation, Formal analysis, Investigation, Data curation, Writing – original draft, Writing – review & editing. Maeva Pinget: Methodology, Investigation, Writing – review & editing. João Sousa: Resources, Methodology, Writing – review & editing. Magda Teles: Conceptualization, Investigation, Writing – review & editing, Supervision, Funding acquisition. Rui F. Oliveira: Conceptualization, Resources, Writing – review & editing, Supervision, Funding acquisition.

## References

1. Calvo, R. & Schluessel, V. Neural substrates involved in the cognitive information processing in teleost fish. Anim Cogn 24, 923–946 (2021).

2. Stednitz, S. J. et al. Forebrain Control of Behaviorally Driven Social Orienting in Zebrafish. Current Biology 28, 2445–2451.e3 (2018).

3. Nunes, A. R. et al. Developmental Effects of Oxytocin Neurons on Social Aailiation and Processing of Social Information. J. Neurosci. 41, 8742–8760 (2021).

4. Pinho, J. S., Cunliffe, V., Kareklas, K., Petri, G. & Oliveira, R. F. Social and asocial learning in zebrafish are encoded by a shared brain network that is differentially modulated by local activation. Commun Biol 6, 633 (2023).

5. Teles, M. C., Almeida, O., Lopes, J. S. & Oliveira, R. F. Social interactions elicit rapid shifts in functional connectivity in the social decision-making network of zebrafish. Proc. R. Soc. B. 282, 20151099 (2015).

6. Cardoso, S. D., Teles, M. C. & Oliveira, R. F. Neurogenomic mechanisms of social plasticity. Journal of Experimental Biology 218, 140–149 (2015).

7. Bludau, A., Royer, M., Meister, G., Neumann, I. D. & Menon, R. Epigenetic Regulation of the Social Brain. Trends in Neurosciences 42, 471–484 (2019).

8. Rustad, S. R., Papale, L. A. & Alisch, R. S. DNA Methylation and Hydroxymethylation and Behavior. in Behavioral Neurogenomics vol. 42 (Springer Berlin Heidelberg, Berlin, Heidelberg, 2019).

9. Danoa, J. S., Connelly, J. J., Morris, J. P. & Perkeybile, A. M. An epigenetic rheostat of experience: DNA methylation of OXTR as a mechanism of early life allostasis. Comprehensive Psychoneuroendocrinology 8, 100098 (2021).

10. Szyf, M. DNA methylation, the early-life social environment and behavioral disorders. J Neurodevelop Disord 3, 238–249 (2011).

11. Jack, A., Connelly, J. J. & Morris, J. P. DNA methylation of the oxytocin receptor gene predicts neural response to ambiguous social stimuli. Front. Hum. Neurosci. 6, (2012).

12. Maud, C., Ryan, J., McIntosh, J. E. & Olsson, C. A. The role of oxytocin receptor gene (OXTR) DNA methylation (DNAm) in human social and emotional functioning: a systematic narrative review. BMC Psychiatry 18, 154 (2018).

13. Puglia, M. H., Lillard, T. S., Morris, J. P. & Connelly, J. J. Epigenetic modification of the oxytocin receptor gene influences the perception of anger and fear in the human brain. Proc. Natl. Acad. Sci. U.S.A. 112, 3308–3313 (2015).

14. Hilliard, A. T., Xie, D., Ma, Z., Snyder, M. P. & Fernald, R. D. Genome-wide effects of social status on DNA methylation in the brain of a cichlid fish, Astatotilapia burtoni. BMC Genomics 20, 699 (2019).

15. Lenkov, K., Lee, M. H., Lenkov, O. D., Swaaord, A. & Fernald, R. D. Epigenetic DNA Methylation Linked to Social Dominance. PLoS ONE 10, e0144750 (2015).

16. Bukhari, S. A. et al. Temporal dynamics of neurogenomic plasticity in response to social interactions in male threespined sticklebacks. PLoS Genet 13, e1006840 (2017).

17. Le Roy, A., Loughland, I. & Seebacher, F. Differential effects of developmental thermal plasticity across three generations of guppies (Poecilia reticulata): canalization and anticipatory matching. Sci Rep 7, 4313 (2017).

18. Bird, A. DNA methylation patterns and epigenetic memory. Genes Dev. 16, 6–21 (2002).

19. Schübeler, D. Function and information content of DNA methylation. Nature 517, 321–326 (2015).

20. Neri, F. et al. Intragenic DNA methylation prevents spurious transcription initiation. Nature 543, 72–77 (2017).

21. Teissandier, A. & Bourc’his, D. Gene body DNA methylation conspires with H3K36me3 to preclude aberrant transcription. The EMBO Journal 36, 1471–1473 (2017).

22. Kriaucionis, S. & Heintz, N. The Nuclear DNA Base 5-Hydroxymethylcytosine Is Present in Purkinje Neurons and the Brain. Science 324, 929–930 (2009).

23. Szulwach, K. E. et al. 5-hmC–mediated epigenetic dynamics during postnatal neurodevelopment and aging. Nat Neurosci 14, 1607–1616 (2011).

24. De Mendoza, A. et al. The emergence of the brain non-CpG methylation system in vertebrates. Nat Ecol Evol 5, 369–378 (2021).

25. Douvlataniotis, K., Bensberg, M., Lentini, A., Gylemo, B. & Nestor, C. E. No evidence for DNA *N*^6^-methyladenine in mammals. Sci. Adv. 6, eaay3335 (2020).

26. O’Brown, Z. K. et al. Sources of artifact in measurements of 6mA and 4mC abundance in eukaryotic genomic DNA. BMC Genomics 20, 445 (2019).

27. Chatterjee, A., Stockwell, P. A., Horsfield, J. A., Morison, I. M. & Nakagawa, S. Base-resolution DNA methylation landscape of zebrafish brain and liver. Genomics Data 2, 342–344 (2014).

28. Rand, A. C. et al. Mapping DNA methylation with high-throughput nanopore sequencing. Nat Methods 14, 411–413 (2017).

29. Simpson, J. T. et al. Detecting DNA cytosine methylation using nanopore sequencing. Nat Methods 14, 407–410 (2017).

30. Liu, X. et al. Unraveling the whole genome DNA methylation profile of zebrafish kidney marrow by Oxford Nanopore sequencing. Sci Data 10, 532 (2023).

31. Kong, Y. et al. Critical assessment of nanopore sequencing for the detection of multiple forms of DNA modifications. Preprint at 10.1101/2024.11.19.624260 (2024).

32. Kareklas, K., Teles, M. C., Nunes, A. R. & Oliveira, R. F. Social zebrafish: *Danio rerio* as an emerging model in social neuroendocrinology. J Neuroendocrinology 35, e13280 (2023).

33. Percie Du Sert, N., et al. The ARRIVE guidelines 2.0: Updated guidelines for reporting animal research. PLoS Biol 18, e3000410 (2020).

